# DNABERT-based explainable lncRNA identification in plant genome assemblies

**DOI:** 10.1101/2022.02.09.479647

**Authors:** Monica F. Danilevicz, Mitchell Gill, Cassandria G. Tay Fernandez, Jakob Petereit, Shriprabha R. Upadhyaya, Jacqueline Batley, Mohammed Bennamoun, David Edwards, Philipp E. Bayer

## Abstract

Long non-coding ribonucleic acids (lncRNAs) have been shown to play an important role in plant gene regulation, involving both epigenetic and transcript regulation. LncRNAs are transcripts longer than 200 nucleotides that are not translated into functional proteins but can be translated into small peptides. Machine learning models have predominantly used transcriptome data with manually defined features to detect lncRNAs, however, they often underrepresent the abundance of lncRNAs and can be biased in their detection. Here we present a study using Natural Language Processing (NLP) models to identify plant lncRNAs from genomic sequences rather than transcriptomic data. The NLP models were trained to predict lncRNAs for seven model and crop species (*Zea mays*, *Arabidopsis thaliana*, *Brassica napus*, *Brassica oleracea*, *Brassica rapa*, *Glycine max* and *Oryza sativa*) using publicly available genomic references. We demonstrated that lncRNAs can be accurately predicted from genomic sequences with the highest accuracy of 83.4% for *Z. mays* and the lowest accuracy of 57.9% for *B. rapa*, revealing that genome assembly quality might affect the accuracy of lncRNA identification. Furthermore, we demonstrated the potential of using NLP models for cross-species prediction with an average of 63.1% accuracy using target species not previously seen by the model. As more species are incorporated into the training datasets, we expect the accuracy to increase, becoming a more reliable tool for uncovering novel lncRNAs. Finally, we show that the models can be interpreted using explainable artificial intelligence to identify motifs important to lncRNA prediction and that these motifs frequently flanked the lncRNA sequence.

**Highlights:**

- We demonstrate for the first time the identification of lncRNAs from genomic sequences, instead of transcriptome sequences, allowing the identification of lowly expressed lncRNAs.
- A deep learning model (natural language processing) was employed to predict lncRNAs in two monocot and five dicot plant species.
- We used explainable machine learning to extract the genomic motifs associated with lncRNA identification and highlighted potentially conserved structures.

## Introduction

Regulatory non-coding ribonucleic acids (ncRNAs) can modulate gene expression, with many ncRNAs reported to regulate flowering time and development [1,2], abiotic stress tolerance [3,4] and interaction beneficial or pathogenic microorganisms [5–7]. Long non-coding ribonucleic acids (lncRNA), a subset of ncRNAs, are defined as RNA transcripts longer than 200 nucleotides that are not translated into functional proteins but can sometimes generate small peptides [8,9]. LncRNA expression is often specific to the plant developmental stage and tissue or cell type [10–13], with some transcripts presenting a remarkably short life [14]. Once considered an evolutionary oddity, lncRNAs have demonstrated roles in essential biological processes. They may act by regulating the expression of neighbouring genes or across multiple loci in the genome, and are also involved in epigenetic regulation [15–17]. For example in rice, the overexpression of lncRNA LAIR can increase grain yield by upregulating the expression of the LRK (leucine-rich repeat receptor kinase) gene cluster, producing larger primary panicles and more panicles per plant [18]. The lncRNA Ef-cd, transcribed from the antisense strand of the flowering activator OsSOC1 locus, positively regulates the expression of this gene, shortening time to maturity in late-maturing rice by 7-20 days, without causing a decrease in yield usually associated with early maturing varieties [19]. Other lncRNAs are associated with heat responsive genes in *Brassica rapa [20]*, osmotic and salt stress in *Medicago truncatula [21]*, drought stress response in *Zea mays* and *Oryza sativa [22,23]*, stress response in *Glycine max [24]*, sexual reproduction in *O. sativa [25]*, pathogen response in *Triticum aestivum* and *Arabidopsis thaliana [26–28]*, and vernalization-mediated epigenetic silencing in *A. thaliana [15]*. Hence the identification of novel lncRNAs may support the breeding of improved crop varieties.

Multiple approaches are employed for lncRNA detection and characterisation. The majority of lncRNA prediction bioinformatics pipelines use a combination of in-house scripts to filter potential lncRNAs from transcriptome datasets based on length, open read frame (ORF) size and comparison to known protein-coding sequences, followed by a tool to estimate coding potential, such as CPC2 [29], LncMachine [30], CPAT [31], PlncPRO [32] and PLEK [33] (reviewed in [34]). For the coding potential models, a common approach is to manually define features to be extracted from transcripts and to use these features as input for machine and deep learning models, such as random forest, to categorise transcripts into mRNA or lncRNA [35,36]. However, handcrafted features used for machine learning can introduce bias and reduce classification performance, depending on the relevance of the chosen features [37,38]. Another drawback is that current models rely on transcriptome data or computationally predicted lncRNAs available on CANTATAdb [39], PLncDB [40], GreeNC [41,42] or AlnC [43] to train their models and output a prediction. Since lncRNAs have a low expression rate that can be highly specific to the plant developmental stage or tissue, identifying lncRNAs from transcriptome datasets may underestimate lncRNA abundance or fail to detect lncRNA presence [24]. A model that could take advantage of publicly available genomic references rather than transcriptomic data to identify lncRNAs would potentially identify all lncRNAs, particularly if trained with learned features, avoiding bias introduced by handcrafted feature extraction.

Natural Language Processing (NLP) models, such as Bidirectional Encoder Representations from Transformers (BERT) [44] and Universal Language Model Fine-tuning (ULMFiT) [45], are deep learning models developed to deal with large sequences of words or text data. Transformers employ an attention-based mechanism to train their encoder-decoder architecture [46], offering an opportunity to initially pre-train a model on genomic data without labelling the dataset beforehand (unsupervised training), avoiding the introduction of annotation bias. DNABERT, GeneBERT and Genomic-ULMFiT are the genomic versions of the NLP frameworks (https://github.com/kheyer/Genomic-ULMFiT) [47,48]. These NLP models have been fine-tuned for the identification of N4-methylcytosine DNA methylation sites in the *Caenorhabditis elegans* genome [49], identification of gene promoters, splice sites and transcription factor binding sites from the human genome data [47], and disease risk estimation based on genomic sequences and RNA-splicing [48]. One of the advantages of using a pre-trained genomic model is that it can be fine-tuned to a specific classification task using a smaller dataset with high quality annotations. NLP offers an opportunity to achieve direct genome prediction of lncRNAs from genomic data. An advantage of generating models from genomic data is that models trained with high accuracy on high quality data will likely have cross-species applicability, with previous evidence in transcriptomics prediction indicating machine learning models trained on well studied crops can predict across species [50]. The NLP models could then be interpreted to determine important motifs in lncRNA prediction within and across plant species to determine evolutionary conserved regulatory motifs. Visualisation and interpretation techniques have been used by DNABERT [47] to pre-train models for genomic input and to visualise important motifs for promoter identification through attention scores [51] for universal applicability to any NLP model that uses transformers. Packages such as these make the leap from predicting promoter regions to lncRNA transcripts feasible with pre-trained BERT language models for interpretation and visualisation.

This study serves as a foundation for performing a genome-wide search and discovery for lncRNAs using an NLP deep learning model. Here, we trained NLP models to predict lncRNA from genomic data of seven model and crop species (*Zea mays, Arabidopsis thaliana, Brassica napus, Brassica oleracea, Brassica rapa, Glycine max and Oryza sativa*) with accuracy between 83.4% and 57.9%. The cross-applicability of these NLP models was tested by comparing each species’ model’s performance with lncRNA data from other species. In addition, we further investigate the model features to visualise the most important motifs for lncRNA prediction. Significant motif conservation across species was also investigated, with the majority of the significant motifs located at the edges of the lncRNA sequence of the species analysed.

## Results

### LncRNA prediction performance of for seven crop species

Multiple NLP model instances were fine-tuned separately for each species using genomic sequences, the sequences were split into different k-mers sizes from 3-mers to 6-mers. The models were evaluated using accuracy, AUROC, F1, MCC, precision and recall. The accuracy reported for the highest performing model in each species ranged from 83.4% and 57.9% (73.5% median). The value for MCC varied substantially, ranging between 0.16-0.67 (0.4 median), with 1 being considered a perfect prediction. AUROC varied between 61% and 90%, while F1 ranged between 58% and 83%, the precision score ranged between 58% and 84% and a recall score between 58% and 84% (Table 1). Across all classifiers, we observed a 30% performance difference, with the *Z. mays* model consistently showing the highest prediction performance, whereas the *B. rapa* model reported the lowest prediction performance. Assessing the impact of k-mers size on model performance, most species achieved the highest performance using the fine-tuned 3-mers model, except for *B. rapa* and *Z. mays,* which had the best accuracy using the 4-mers and 6-mers models respectively.

**Table 1.** The performance metrics of different k-mers length NLP models are fine-tuned for each species. The best metric model for each species is presented in bold text.

| Species | K-mer | Accuracy | AUROC | F1 | MCC | Precision | Recall |
| --- | --- | --- | --- | --- | --- | --- | --- |
| *Arabidopsis thaliana* | **3** | **65.39** | **0.72** | **0.65** | **0.31** | **0.65** | **0.65** |
|  | 4 | 64.82 | 0.72 | 0.65 | 0.30 | 0.65 | 0.65 |
|  | 5 | 63.30 | 0.69 | 0.63 | 0.27 | 0.63 | 0.63 |
|  | 6 | 63.88 | 0.71 | 0.64 | 0.28 | 0.64 | 0.64 |
| *Glycine max* | **3** | **72.77** | **0.79** | **0.73** | **0.45** | **0.73** | **0.73** |
|  | 4 | 71.62 | 0.8 | 0.71 | 0.44 | 0.72 | 0.72 |
|  | 5 | 71.62 | 0.80 | 0.71 | 0.43 | 0.72 | 0.72 |
|  | 6 | 71.39 | 0.80 | 0.71 | 0.43 | 0.72 | 0.71 |
| *Brassica napus* | **3** | **74.6** | **0.81** | **0.74** | **0.49** | **0.75** | **0.74** |
|  | 4 | 71.9 | 0.80 | 0.70 | 0.44 | 0.73 | 0.71 |
|  | 5 | 71.9 | 0.80 | 0.71 | 0.44 | 0.73 | 0.71 |
|  | 6 | 72.4 | 0.80 | 0.71 | 0.45 | 0.73 | 0.71 |
| *Brassica oleracea* | **3** | **74.15** | **0.81** | **0.74** | **0.49** | **0.75** | **0.74** |
|  | 4 | 72.9 | 0.80 | 0.73 | 0.46 | 0.729 | 0.73 |
|  | 5 | 70.8 | 0.79 | 0.71 | 0.41 | 0.708 | 0.71 |
|  | 6 | 73.1 | 0.80 | 0.72 | 0.46 | 0.74 | 0.72 |
| *Brassica rapa* | 3 | 57.7 | **0.63** | 0.58 | **0.16** | **0.58** | **0.58** |
|  | **4** | **57.86** | 0.61 | **0.58** | 0.16 | 0.58 | 0.58 |
|  | 5 | 56.86 | 0.60 | 0.58 | 0.14 | 0.57 | 0.57 |
|  | 6 | 56.49 | 0.60 | 0.56 | 0.13 | 0.57 | 0.57 |
| *Oryza sativa* | **3** | **61.65** | **0.65** | **0.62** | **0.23** | **0.62** | **0.62** |
|  | 4 | 61.11 | 0.65 | 0.61 | 0.22 | 0.61 | 0.61 |
|  | 5 | 58.75 | 0.63 | 0.59 | 0.18 | 0.59 | 0.59 |
|  | 6 | 58.86 | 0.63 | 0.59 | 0.18 | 0.59 | 0.59 |
| *Zea mays* | 3 | 81.2 | 0.90 | 0.81 | 0.64 | 0.83 | 0.81 |
|  | 4 | 83.1 | 0.91 | 0.83 | 0.67 | 0.83 | 0.83 |
|  | 5 | 81.94 | 0.90 | 0.82 | 0.65 | 0.83 | 0.82 |
|  | **6** | **83.42** | **0.90** | **0.83** | **0.67** | **0.84** | **0.84** |

Confusion matrix analysis is commonly used in machine learning classification tasks to assess which classes are most often misclassified (or “confused”) by the model, it helps identify whether the misclassified samples share similarities and if either class is more often wrongly predicted. Here, the models accurately identified lncRNAs 59.3 to 88.6% of the time and detected random stretches of DNA between 55.6 to 78.1% of the time, showing the model was equally accurate in predicting either class. The *Z. mays* model had the largest dataset with 14030 genomic sequences and achieved the highest classification performance to identify both lncRNAs and no lncRNAs. The two models trained with *G. max* and *O. sativa* had smaller datasets with 3414 and 3716 sequences, respectively, but performed on average 18.7% better than the *B. rapa* model with a medium-size set (9725 genomic sequences).

### Identification of lncRNA from external datasets across six species

Machine learning models are often impacted by lack of variability within a training dataset, which might lead to lower performance when faced with external data. In this section, we evaluated the robustness of the proposed methodology to detect lncRNAs across seven datasets dedicated to lncRNA curation. Including a wide-range of lncRNAs databases in the model evaluation phase challenges to the model’s capacity to generalize and sustain consistent performance across a spectrum of conditions, demonstrating its capacity to classify transcripts extracted from multiple sequencing experiments and a different genome sequence than the one employed for constructing the training and validation dataset in the previous section.

In total, this section encompasses 21,491 lncRNAs sourced from PLncDB (7,795), RNAcentral (1,747), EVLncRNAs (55), Liu (4,501), NCBI (2,688), Li (2,392), and Cameron (2,313). These lncRNAs were predicted using best performing fine-tuned models established in the previous section. The comprehensive evaluation of prediction performance across six species shows that 65% of the lncRNAs were accurately classified. The models exhibited a consistent 65% for both Recall and Precision with an average F1 score of 0.78, which is compatible with the results observed in the previous section.

A substantial variation in prediction performance is observed across the species as shown in Table 2. The highest prediction performance is observed in the identification of transcripts from *O. sativa* and *G. max* models, which showed 67.4% and 76.3% accuracy respectively. Both species models outperformed the F1 score by at least 0.10 in comparison to the previous performance observed with the Cantata dataset. The *Z. mays* prediction model showed the lowest classification performance observed, correctly classifying 54% of the lncRNAs which is 30% less than the lncRNA prediction accuracy on the Cantata database. Moreover, the *B. napus* specific model presented a 16% drop in prediction accuracy, precision and recall for the identification of lncRNAs in the external datasets in comparison to the results obtained with Cantata. There were 4,563 transcripts identified as long intergenic non-coding RNAs (lincRNAs) with the majority of these being from *A. thaliana* (4,536) and the remaining from *O. sativa* (14), *G. max* (8) and *Z. mays* (5). The prediction performance of lincRNAs showed a similar performance to the species specific models described in Table 2. A detailed list of the lncRNA sequence, prediction label, chromosomal position and attributes is available in Supplementary Table 1.

**Table 2:** Evaluation metrics of species specific models using the best fine-tuned models presented in the previous section for the identification of lncRNAs from PLncDB, RNAcentral, EVLncRNAs, Liu, NCBI, Li, and Cameron datasets.

| Species | K-mer | Accuracy | F1 | Precision | Recall |
| --- | --- | --- | --- | --- | --- |
| *Arabidopsis thaliana* | 3 | 63.68 | 0.78 | 0.64 | 0.64 |
| *Glycine max* | 3 | 76.31 | 0.87 | 0.76 | 0.76 |
| *Brassica napus* | 3 | 58.63 | 0.74 | 0.59 | 0.59 |
| *Brassica rapa* | 4 | 57.83 | 0.74 | 0.59 | 0.59 |
| *Oryza sativa* | 3 | 67.45 | 0.81 | 0.67 | 0.67 |
| *Zea mays* | 6 | 54.50 | 0.71 | 0.55 | 0.55 |

### Cross-species prediction of lncRNAs using the single species fine-tuned NLP model

NLP models with the highest accuracy for each crop were used to predict the lncRNAs in another species. For example, the 3-mers model trained with *A. thaliana* genomic k-mers was employed to predict lncRNAs in the remaining six species without further fine-tuning. Most of the models performed similarly or slightly worse at predicting lncRNAs for a new species in comparison to identifying lncRNAs in their original species, except for the model trained on *Z. mays* data that performed significantly worse at classifying other species’ lncRNAs (Table 3, Figure 2). In all cases, the highest lncRNA classification performance was achieved by models trained in that species instead of cross-species prediction models.

**Figure 1:**
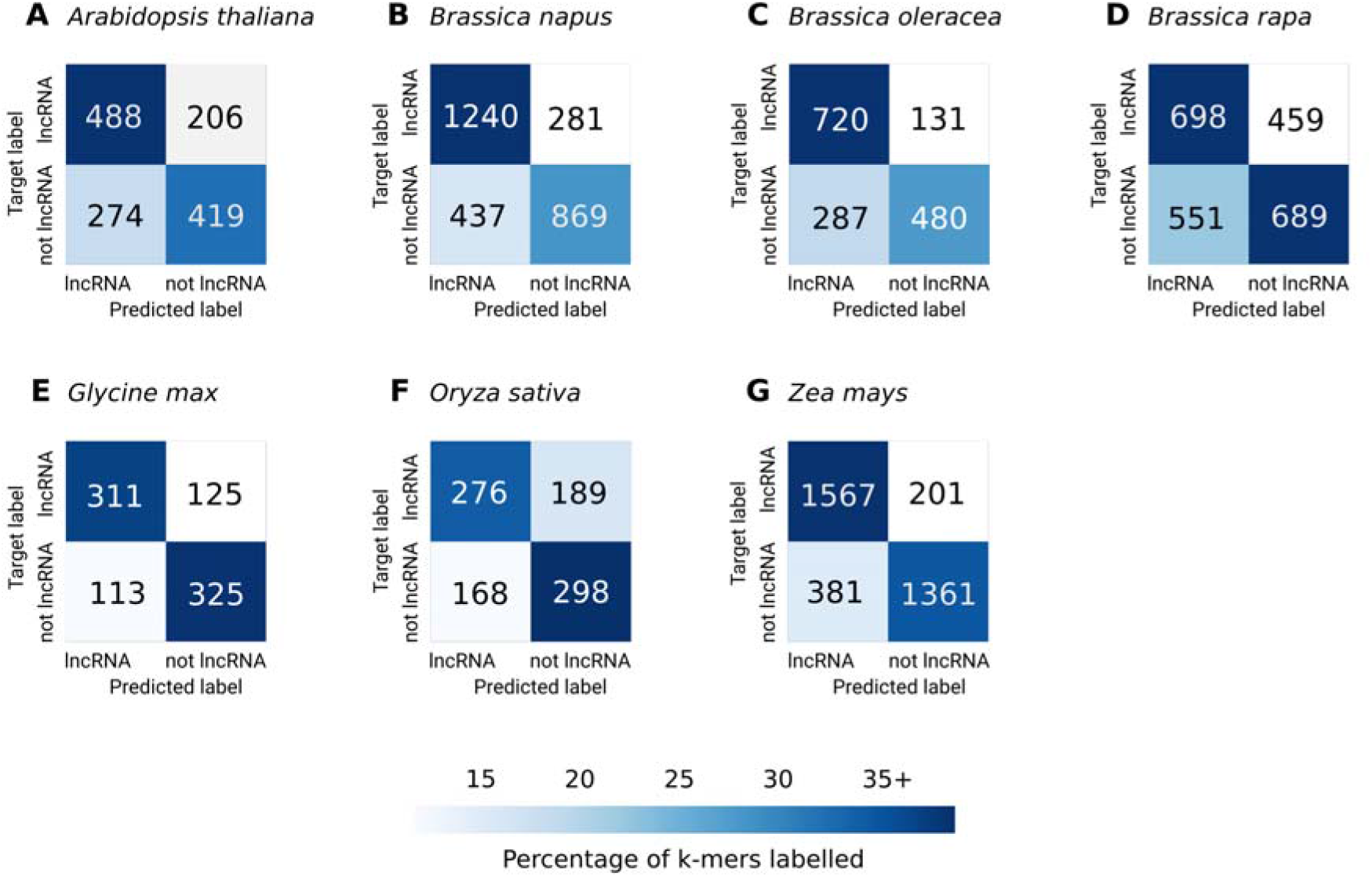
Confusion matrices of the predicted labels from each species’ highest performing models. The matrix is coloured based on the percentage of k-mers labelled in each class from the total test set of the species indicated in the title (i.e. *A. thaliana* had 1387 k-mers in total, with 488 of these (above 35%) being correctly predicted as lncRNAs)

**Figure 2:**
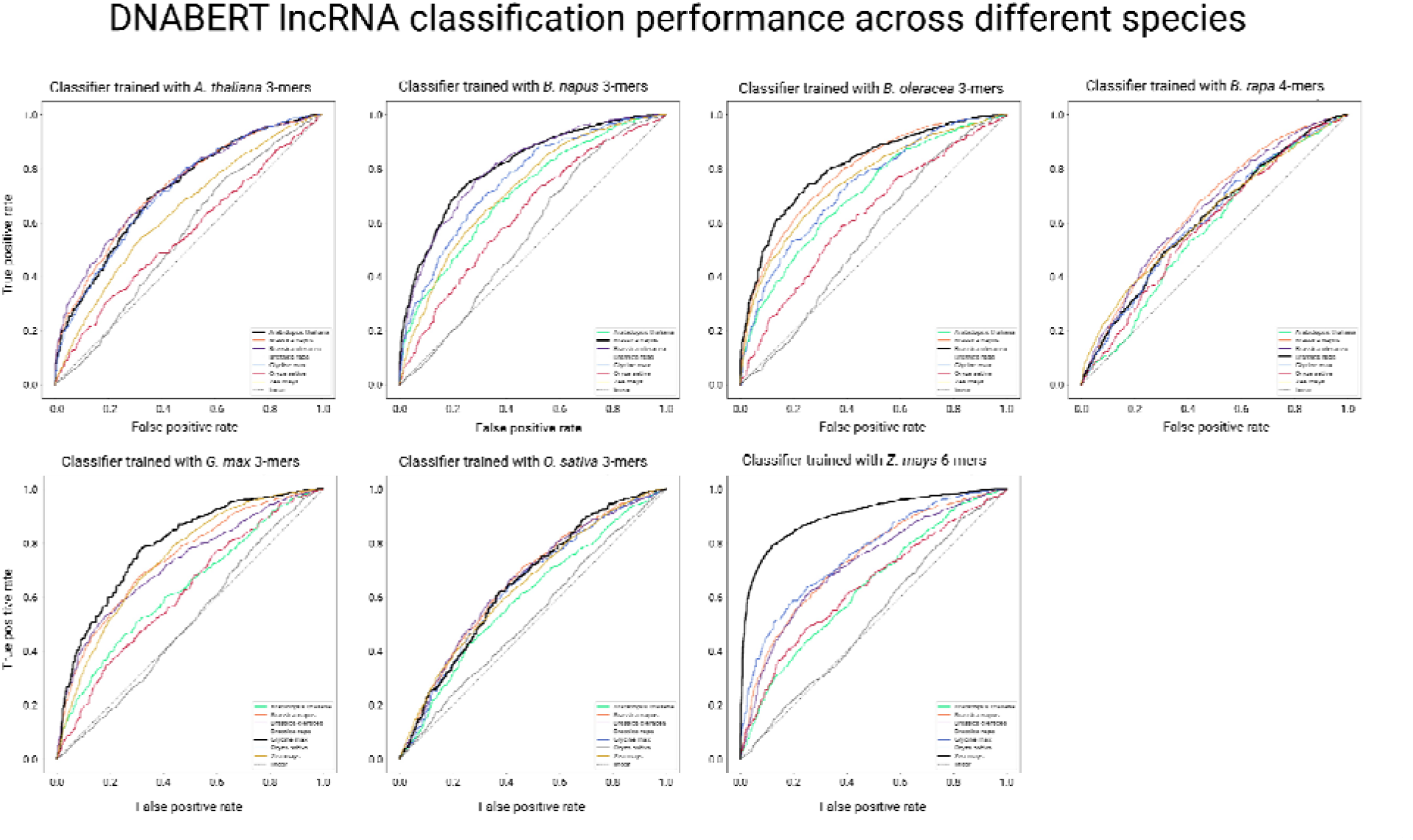
AUROC of each fine-tuned model when performing cross-species prediction of lncRNAs. The black line indicates model performance when predicting lncRNAs in the species it was trained, whereas other species are represented by the colours indicated in the plot legend.

**Table 3:** Performance metrics of cross-species lncRNA prediction.

| Trained Species | Test Species | K-mers | Accuracy | AUROC | F1 | MCC | Precision | Recall |
| --- | --- | --- | --- | --- | --- | --- | --- | --- |
| *A. thaliana* | *B. napus* | 3 | 0.67 | 0.73 | 0.67 | 0.34 | 0.67 | 0.67 |
|  | *B. oleracea* | 3 | 0.67 | 0.73 | 0.67 | 0.33 | 0.67 | 0.66 |
|  | *B. rapa* | 3 | 0.56 | 0.55 | 0.55 | 0.11 | 0.55 | 0.55 |
|  | *G. max* | 3 | 0.65 | 0.72 | 0.65 | 0.30 | 0.65 | 0.65 |
|  | *O. sativa* | 3 | 0.53 | 0.56 | 0.53 | 0.06 | 0.53 | 0.53 |
|  | *Z. mays* | 3 | 0.58 | 0.64 | 0.57 | 0.18 | 0.60 | 0.58 |
| *B. napus* | *A. thaliana* | 3 | 0.65 | 0.71 | 0.65 | 0.30 | 0.65 | 0.65 |
|  | *B. oleracea* | 3 | 0.72 | 0.81 | 0.72 | 0.45 | 0.73 | 0.72 |
|  | *B. rapa* | 3 | 0.54 | 0.56 | 0.54 | 0.08 | 0.54 | 0.54 |
|  | *G. max* | 3 | 0.69 | 0.76 | 0.69 | 0.38 | 0.69 | 0.69 |
|  | *O. sativa* | 3 | 0.58 | 0.63 | 0.58 | 0.17 | 0.59 | 0.58 |
|  | *Z. mays* | 3 | 0.66 | 0.72 | 0.65 | 0.32 | 0.66 | 0.66 |
| *B. oleracea* | *A. thaliana* | 3 | 0.64 | 0.71 | 0.63 | 0.28 | 0.65 | 0.64 |
|  | *B. napus* | 3 | 0.71 | 0.79 | 0.7 | 0.41 | 0.71 | 0.7 |
|  | *B. rapa* | 3 | 0.53 | 0.55 | 0.53 | 0.07 | 0.53 | 0.53 |
|  | *G. max* | 3 | 0.66 | 0.73 | 0.65 | 0.32 | 0.66 | 0.66 |
|  | *O. sativa* | 3 | 0.59 | 0.62 | 0.58 | 0.18 | 0.59 | 0.59 |
|  | *Z. mays* | 3 | 0.69 | 0.75 | 0.69 | 0.38 | 0.69 | 0.69 |
| *B. rapa* | *A. thaliana* | 4 | 0.55 | 0.57 | 0.55 | 0.10 | 0.55 | 0.55 |
|  | *B. napus* | 4 | 0.61 | 0.65 | 0.61 | 0.21 | 0.61 | 0.60 |
|  | *B. oleracea* | 4 | 0.61 | 0.65 | 0.61 | 0.22 | 0.61 | 0.61 |
|  | *G. max* | 4 | 0.59 | 0.61 | 0.59 | 0.18 | 0.59 | 0.59 |
|  | *O. sativa* | 4 | 0.57 | 0.59 | 0.57 | 0.14 | 0.57 | 0.57 |
|  | *Z. mays* | 4 | 0.58 | 0.62 | 0.58 | 0.16 | 0.58 | 0.58 |
| *G. max* | *A. thaliana* | 3 | 0.59 | 0.63 | 0.59 | 0.19 | 0.59 | 0.59 |
|  | *B. napus* | 3 | 0.68 | 0.74 | 0.68 | 0.37 | 0.68 | 0.68 |
|  | *B. oleracea* | 3 | 0.66 | 0.73 | 0.66 | 0.32 | 0.66 | 0.66 |
|  | *B. rapa* | 3 | 0.51 | 0.51 | 0.50 | 0.09 | 0.50 | 0.50 |
|  | *O. sativa* | 3 | 0.58 | 0.62 | 0.57 | 0.16 | 0.58 | 0.58 |
|  | *Z. mays* | 3 | 0.67 | 0.74 | 0.67 | 0.34 | 0.67 | 0.67 |
| *O. sativa* | *A. thaliana* | 3 | 0.56 | 0.60 | 0.55 | 0.13 | 0.57 | 0.56 |
|  | *B. napus* | 3 | 0.60 | 0.66 | 0.60 | 0.23 | 0.62 | 0.61 |
|  | *B. oleracea* | 3 | 0.60 | 0.66 | 0.59 | 0.22 | 0.62 | 0.60 |
|  | *B. rapa* | 3 | 0.53 | 0.53 | 0.50 | 0.05 | 0.53 | 0.52 |
|  | *G. max* | 3 | 0.60 | 0.64 | 0.59 | 0.20 | 0.61 | 0.60 |
|  | *Z. mays* | 3 | 0.59 | 0.64 | 0.58 | 0.19 | 0.60 | 0.59 |
| *Z. mays* | *A. thaliana* | 6 | 0.59 | 0.64 | 0.58 | 0.20 | 0.60 | 0.59 |
|  | *B. napus* | 6 | 0.68 | 0.74 | 0.67 | 0.35 | 0.68 | 0.67 |
|  | *B. oleracea* | 6 | 0.68 | 0.72 | 0.67 | 0.35 | 0.68 | 0.67 |
|  | *B. rapa* | 6 | 0.50 | 0.52 | 0.50 | 0.00 | 0.50 | 0.50 |
|  | *G. max* | 6 | 0.69 | 0.76 | 0.69 | 0.39 | 0.70 | 0.69 |
|  | *O. sativa* | 6 | 0.61 | 0.64 | 0.59 | 0.25 | 0.64 | 0.61 |

In addition, we assessed how a model trained on a monocot (*Z. mays* and *O. sativa*) or dicot (*A. thaliana*, *B. rapa*, *B. oleracea*, *B. napus* and *G. max*) species would perform when predicting lncRNAs from the other group. Here, the models performed better when predicting lncRNAs from the same group, with an average accuracy of 66.2% for monocot-monocot prediction, and 62.7% for dicot-dicot prediction. Models performing lncRNA prediction in a different group (monocot-dicot and dicot-monocot) presented an average accuracy of 60.3%.

### Motif identification and frequency of motifs appearing per genomic location

The characterisation of lncRNAs is a challenging task as lncRNAs share many similarities with other RNA types. Thus, to uncover potentially conserved patterns in lncRNA sequences, we extracted and ranked the most relevant genomic features employed by the best performing models for the classification of lncRNA. The best performing model for each crop (ranked by accuracy) was interpreted using the transformers interpret package [51] using the lncRNA sequences from the holdout dataset. The 50 most important genomic motifs for lncRNA identification were subjected to a chi-squared test against non-lncRNA motifs to determine the significant motifs (p-value < 0.05). The resulting 158 unique motifs were compared across species according to sequence similarity (Figure 2, Supplementary Table 2-3). Apart from *O. sativa*, which shared no significant motifs with any of the other species, all species shared two lncRNA motifs (AAAAAAA and AAAAAAAA) belonging to the same motif cluster. The *Brassica* species shared five motifs, whereas *B. napus* and *B. oleracea* shared twelve motifs. *Z. mays* and *G. max* shared eleven motifs, the second highest number shared between two species. Visualising the location of significant motifs in lncRNA divided into 25 bp bins of upstream & downstream flanking regions and bins representing 5% lncRNA sequence, showed that significant motifs important to lncRNA appeared more frequently in the upstream and downstream flanking regions than in the lncRNA sequences themselves (Figure 4, Supplementary Figure 1).

**Figure 3:**
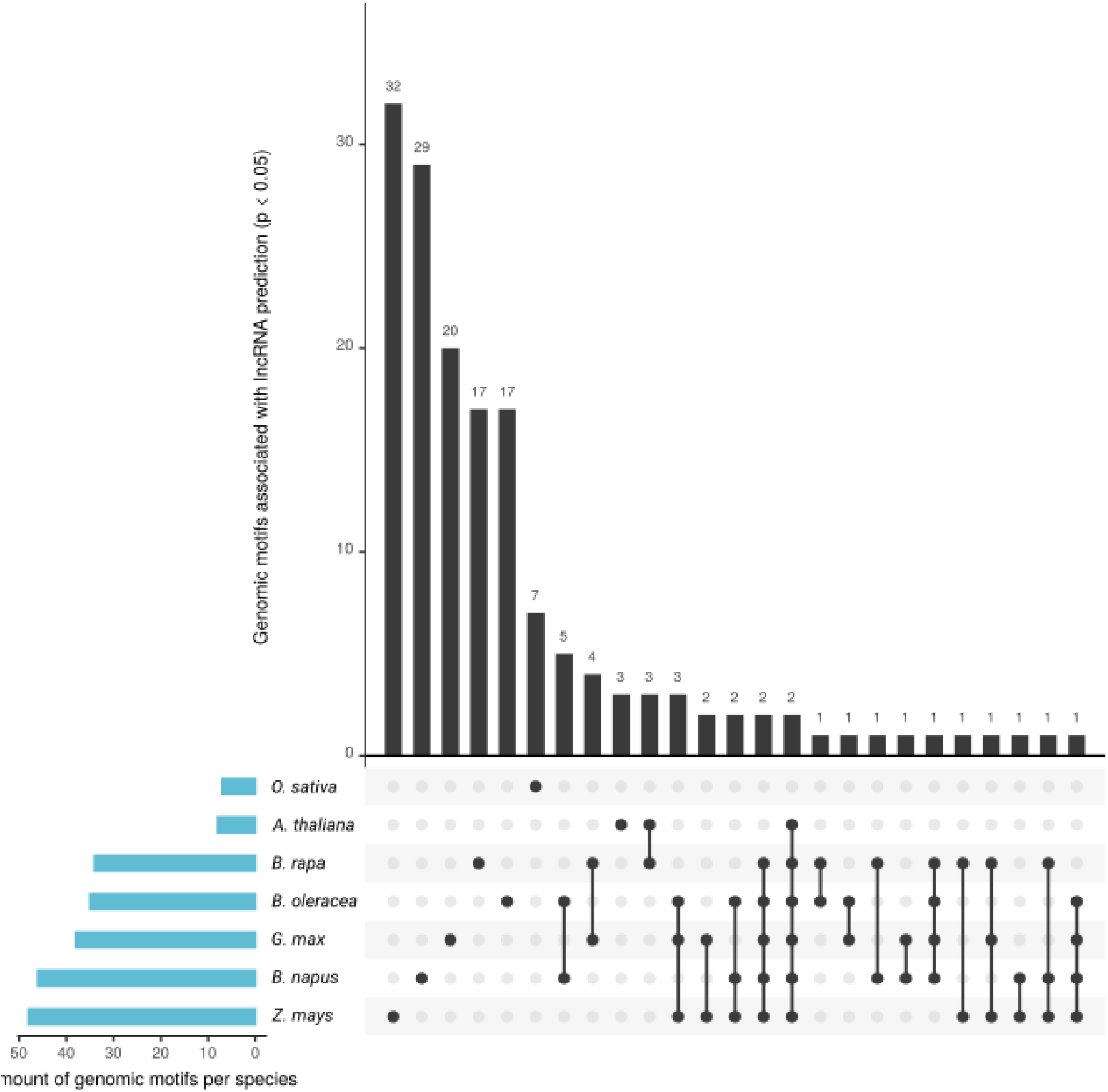
Shared genomic motifs across different plant species. The main plot indicates the number of significant genomic motifs (p-value < 0.05) shared or unique to the species. The number of significant genomic motifs identified per species is indicated in the lateral plot in blue.

**Figure 4:**
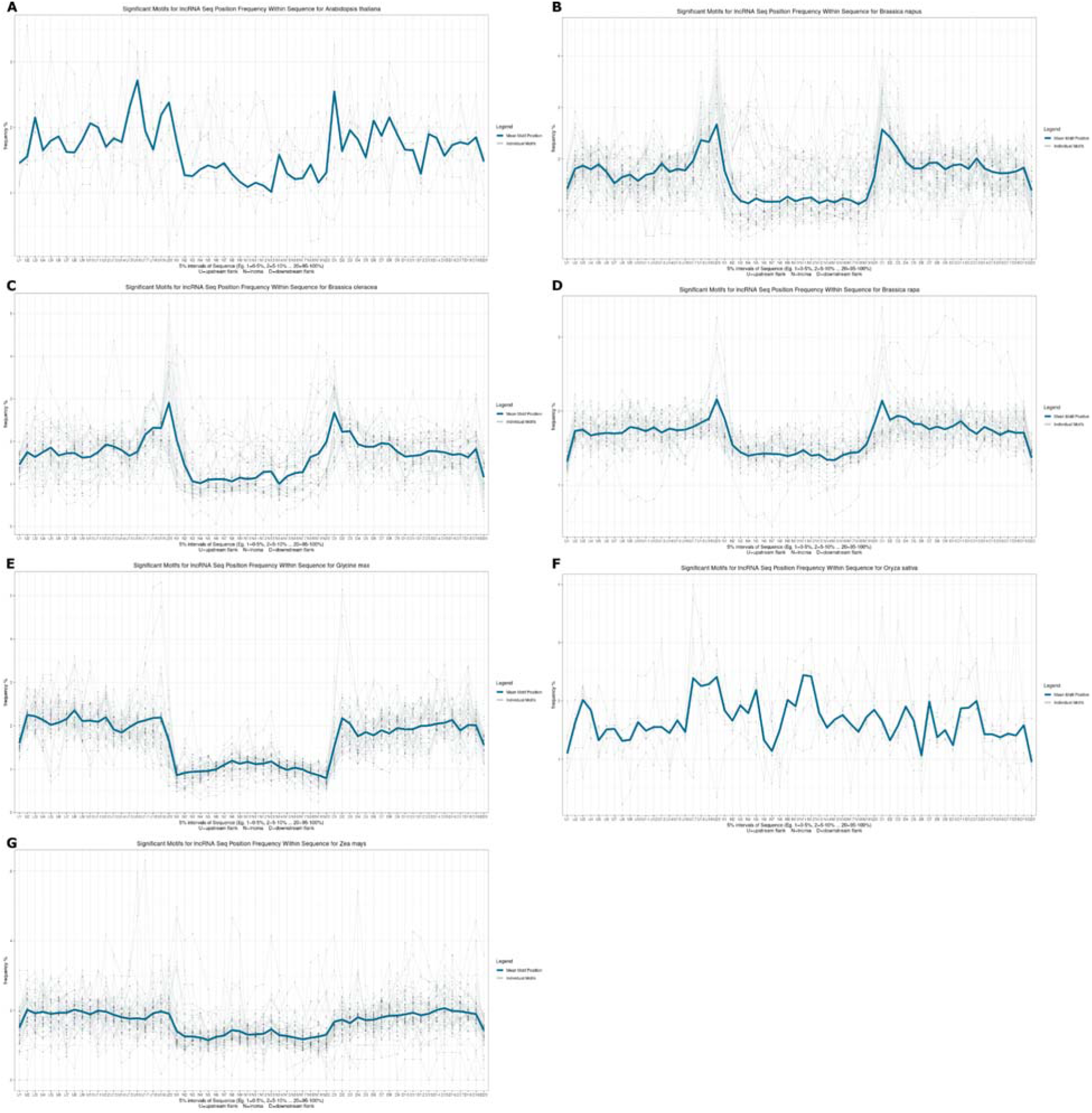
Significant (p-value < 0.05) genomic motif sequence location for identifying lncRNAs in each species based on the highest performance single species classification model.

## Discussion

In this study, we fine-tuned multiple NLP instances to successfully identify lncRNAs within the genome sequence of seven plant species without the need for transcriptome data, and used these models for cross-species prediction of lncRNAs. The code used to develop this project is available on Github (https://github.com/AppliedBioinformatics/lncRNA_Prediction_Interpretation) along with the trained model weights, allowing other researchers to implement the model with their datasets. Identifying lncRNA transcripts is challenging as lncRNAs tend to have lower expression than other transcripts, and their expression is often specific to the development stage, tissue or cell type [10–13] causing lncRNAs to go undetected when using RNA-seq based models [12]. To address this limitation, we identified lncRNAs directly from the genomic sequence using known lncRNAs and genome assemblies. Identifying lncRNAs from genomic sequences can further our understanding of their sequence conservation across plant species [24], providing a foundation to investigate lncRNA evolution and identify novel transcripts [52]. The NLP models fine-tuned per species presented a median accuracy of 73.5%, with the model trained with *Z. mays* performing 25.6% better than the model for *B. rapa*. Several factors may have contributed to the higher performance of the *Z. mays* model, including the high quality of genome assembly. The *Z. mays* AGP RefGen 4v genome assembly was constructed using advanced single-molecule technology, improving contig length and supporting the accurate assembly of intergenic spaces and centromeres [53], providing higher quality sequences for model training. Centromeric regions are reported to encode actively transcribed lncRNAs [24] associated with centromere maintenance and cellular division [54]. In our study, *Z. mays* also had the largest available dataset of predicted lncRNAs after filtering [55], which supports the accurate training of machine learning models as they may better represent transcript diversity [56]. The lncRNAs from *B. rapa* and *O. sativa* were consistently misclassified across the different models, which may indicate that different features were more important for these species. The difficulty in accurately identifying lncRNAs from these species may also be associated with genome assembly and annotation quality. *B. rapa* was the first species sequenced among the *Brassica* genus in 2011 [57], and the *O. sativa* genome was annotated before RNA sequencing was widely available, so these genome assemblies and annotations might be missing lncRNA regions [25,58]. It was previously reported in the *O. sativa* genome annotation that many lncRNA loci had been incorrectly classed as putative protein-coding genes [41]. Further research will measure the impact of genome assembly on model performance to assess the importance of assembly methods on lncRNA identification. The algorithm architecture applied here is susceptible to dataset errors, as it learns directly from the data presented, thus re-training the model as more validated lncRNA sequences are released will lead to higher model accuracy. Since the model presented here is trained using known lncRNAs identified through bioinformatics pipelines, it is difficult to assess its ability to identify novel lncRNAs, though the predicted set may provide a foundation for subsequent validation.

Although lncRNAs do not show the same degree of interspecies sequence conservation as protein-coding genes [59], a study using ten different monocot and dicot species showed that the majority of lncRNAs have some sequence conservation at the intra- and sub-species level, and the number of lncRNAs identified correlated with the number of genes in each species [60]. There is also evidence of structural and positional conservation across animal lncRNAs showing similar expression patterns suggesting transcript functional conservation [61,62]. In our study, the models trained on monocot or dicot species had, on average, 3-6% higher accuracy than when predicting lncRNAs from other species in the same group, suggesting differences in the features employed for predicting these groups. The trained NLP models identified multiple genomic motifs significantly associated with lncRNA prediction. For the majority of species, the genomic motifs for lncRNA identification were more frequently located at the boundaries of the lncRNA sequence (Figure 3), and presented a low GC content in comparison to the lncRNA sequences, with several being composed of poly-A or poly-T stretches. Characterising the role of these conserved motifs in lncRNA expression is beyond the scope of this study and the datasets, however, these motifs may contribute to identifying lncRNA expression regulation in future research. In addition, as more conserved motifs are identified, these may act as novel genomic markers allowing more precise identification of lncRNAs from other plant species.

## Conclusion

We present the first study using an NLP model to successfully identify lncRNAs in plant species from genomic sequences instead of transcriptomic data. This method might allow for the detection of lncRNAs from extracted directly from the genome reference regardless of their expression and can contribute to our understanding of lncRNA conservation and function across plant species. The models developed here are able to identify the majority of lncRNAs from the species used for training, and across different species, with the *Z. mays* model presenting the highest accuracy. Further research may include a wider diversity of genomic sequences to increase model robustness and achieve better classification performance in the genome-wide identification of lncRNAs.

## Methods

### Data Acquisition

The lncRNA datasets from *Arabidopsis thaliana, Brassica napus, Brassica oleracea, Brassica rapa, Glycine max, Oryza sativa* and *Zea mays* were downloaded from cantataDB2.0 (Szcześniak et al., 2019). External datasets for model validation were sourced from PLncDB 2.0 [40] for six plant species (*Arabidopsis thaliana, Brassica napus, Brassica oleracea, Brassica rapa, Glycine max, Oryza sativa* and *Zea mays)* as *B. oleracea* was not available. The PLncDB 2.0 database downloaded included 21,491 transcripts from PLncDB [40], RNAcentral [63,64], NCBI [65] and EVLncRNAs 2.0 that hosts experimentally validated lncRNAs [66]. Reference genomes were obtained from Ensembl Plants (release 56) for each of the above crops matching the version used by the CANTATA to extract the corresponding lncRNA genomic sequence. These were as follows; Arabidopsis_thaliana.TAIR10.dna.toplevel.fa, Brassica_napus.AST_PRJEB5043_v1.dna.toplevel.fa, Brassica_oleracea.v2.1.dna.toplevel.fa, Brassica_rapa.IVFCAASv1.dna.toplevel.fa, Glycine_max.V1.0.dna.toplevel.fa, Oryza_sativa.IRGSP-1.0.dna.toplevel.fa and Zea_mays.AGPv4.dna.toplevel.fa. Alternative genome references were obtained from NCBI to match PLncDB 2.0 lncRNAs, including *B. napus* (GCF_000686985.2), *B. rapa* (GCF_000309985.2), *G. max* (GCF_000004515.5), *Z. mays* (Zm-B73-REFERENCE-GRAMENE-4.0)

### BlastX Filtering

The lncRNAs fasta sequences were extracted using bedtools getfasta query [67] using the reference genome fasta for the respective species. A BLAST database for each species was built using makeblastdb query and the genome fasta file [68] Next, the lncRNA fasta sequences for each species was compared to the respective genome database through blastn (outfmt 6, evalue 1e-10) to filter out sequences with high similarity to known mRNAs. The extract_blast.py python script was used to extract lncRNAs with over 90% of its length matched to a mRNA sequence. The blastx_pipeling.sh script can be used to complete these steps efficiently.

### Adding Flanking sequence & Generating a Control Set

The lncRNA genomic context was incorporated in the sequence using the add_flanks.py script, which adds the 500bp flanking regions upstream and downstream of the gtf file. The script ensured the flanking regions did not exceed chromosome or scaffold boundaries using the index file generated with samtools faidx [69]. The control dataset, labelled “non-lncRNA”, was composed from random stretches of genomic. After filtering the lncRNA plus flanking regions from the genome file using bedtools shuffle (-excl lncRNA gtf file), the control sequences were extracted from the reference genome running bedtools getfasta with random intervals of equivalent length to the flanked lncRNA fasta sequences.

### Fine-tuning of DNABERT Models

The 3-mer, 4-mer, 5-mer and 6-mer pretrained BERT models for DNA sequences (DNABERT) were cloned from the DNABERT [47] repository (https://github.com/jerryji1993/DNABERT). The models were trained on GPU servers using a container environment using a tensorflow image (version ‘tensorflow:20.03-tf2-py3’) through singularity [70]. Jupyter notebooks, custom scrips and processed datasets are available at https://github.com/AppliedBioinformatics/lncRNA_Prediction_Interpretation.

Packages were installed as described in the DNABERT repository with the exception of pytorch (Paszke et al., 2019) version 1.4 and Biopython version 1.78 (Cock et al., 2009). Functions from Genomic-ULMFiT (https://github.com/kheyer/Genomic-ULMFiT) were used to parse fasta files and split datasets into training and testing partitions. The input sequences were converted to k-mers using the ‘seq2k-mer’ function from DNABERT.

The DNABERT models were fine-tuned using run_fine-tune.py with the following settings; model_type=dnalongcat, tokenizer_name=dna<K-mers of model>, task_name=dnaprom,-- do_train, --do_eval, max_seq_length=2048, per_gpu_eval_batch_size=4, per_gpu_train_batch_size=4, learning_rate=2e-5, num_train_epochs=4.0, -- evaluate_during_training, logging_steps=100, save_steps=4000, warmup_percent=0.1, hidden_dropout_prob=0.1, --overwrite_output, weight_decay=0.01, n_process=8. The resulting fine-tuned model was evaluated on the separate testing set.

For cross species prediction, the best performing k-mers model for each crop was used to predict the separate test set of every other plant with the same evaluation metrics..

### Evaluation metrics

Model performance was assessed using accuracy, area under receiver operating characteristics curve (AUROC), F-score (F1), Mathew’s correlation coefficient (MCC), Precision and Recall as the evaluation metrics calculated using the scikit-learn library [71]. AUROC is estimated by plotting true positive rate vs false positive rate. The other metrics are defined below, where TP, FP, TN and TP mean true positive, false positive, true negative and true positive:

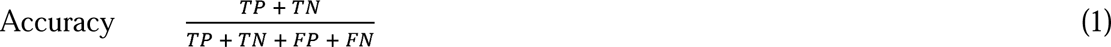

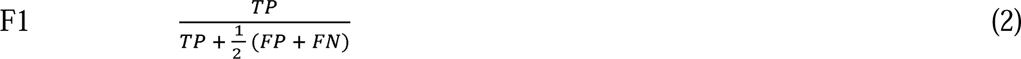

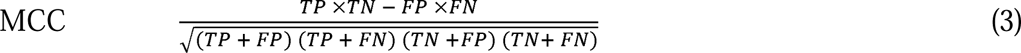

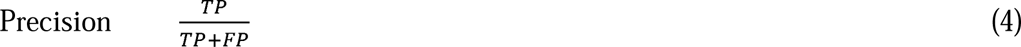

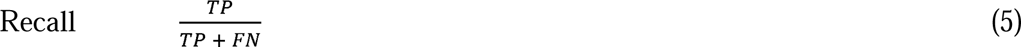

### Interpretation of Models

The transformers interpret repository (https://github.com/cdpierse/transformers-interpret) was used to analyse the model classification results. The scripts atributions.py and explainer.py from this repository were adapted to increase the maximum sequence length of the input from 512 to 2048.

Each lncRNA labelled sequence from the test dataset was iterated over with the transformers interpret sequence classification explainer object. This object returned attribution scores for each k-mer in the sequence. If more than 7 successive k-mers had positive attribution scores for lncRNA prediction, these k-mers were merged and their attribution scores summed until a k-mer with a negative attribution score was present. This merged k-mer would be annotated as a motif important for lncRNA prediction. After iterating over every lncRNA sequence for a given plant species, the frequency of a motif occurring as a positive prediction factor was multiplied by the sum of a motif’s attribution score to give a ranking of the motif’s importance to lncRNA prediction for each DNABERT model. The top 50 motifs were extracted for visualisation.

### Visualisation of Significant Motif Location

The motifs contained in the lncRNA (plus flanking regions) and non-lncRNA sequences were extracted and counted using grep and awk commands. Then, a custom R script (chi_script.R) was used to run a chi-square test to assess whether the motif appeared in lncRNA sequences significantly more than non-lncRNA sequences. Motifs with a p-value less than 0.05 were kept whereas every other motif was excluded.

The parse_fasta function from Genomic_ULMFiT was used to count the locations of significant motifs in lncRNA and non-lncRNA sequences. Each lnRNA sequence was divided into 60 bins. Bins 1-20 were 5% intervals of the upstream flanking region, bins 21-40 were 5% intervals of the lncRNA sequence, and bins 41-60 were 5% intervals of the downstream flanking region. The counts bins 21-40 that refer to the lncRNA sequence, were normalised to counts of bins 1-20 and bins 41-60, as the lncRNA sequence was of variable length whereas the flanks were a consistent 500bp. This ensured motif counts on longer lncRNAs sequences were not biased by the sequence length. The bins and their counts were analysed with the tidyverse suite [72] and plotted with ggplot2 [73] in R.

## Acknowledgments

This work was supported by resources provided by the Pawsey Supercomputing Research Centre with funding from the Australian Government and the Government of Western Australia. The Australian Government supported this work through the Australian Research Council (Projects DP210100296, DP200100762, and DE210100398) and the Grains Research and Development Corporation (Projects 9177539 and 9177591). Monica F. Danilevicz, Mitchell Gill and Cassandria G. Tay Fernandez were supported by the Research Training Program scholarship. Shriprabha Upadhyaya was supported by the Ad Hoc Postgraduate Scholarship. The Forrest Research Foundation further supported Monica F. Danilevicz.

## Bibliography

[1] Lee H, Yoo SJ, Lee JH, Kim W, Yoo SK, Fitzgerald H, et al. Genetic framework for flowering-time regulation by ambient temperature-responsive miRNAs in Arabidopsis. Nucleic Acids Res 2010;38:3081–93. 10.1093/nar/gkp1240.

[2] Waheed S, Zeng L. The Critical Role of miRNAs in Regulation of Flowering Time and Flower Development. Genes (Basel) 2020;11. 10.3390/genes11030319.

[3] Thiebaut F, Rojas CA, Almeida KL, Grativol C, Domiciano GC, Lamb CRC, et al. Regulation of miR319 during cold stress in sugarcane. Plant Cell Environ 2012;35:502–12. 10.1111/j.1365-3040.2011.02430.x.

[4] Liu Q, Yang T, Yu T, Zhang S, Mao X, Zhao J, et al. Integrating Small RNA Sequencing with QTL Mapping for Identification of miRNAs and Their Target Genes Associated with Heat Tolerance at the Flowering Stage in Rice. Front Plant Sci 2017;8:43. 10.3389/fpls.2017.00043.

[5] Hu G, Hao M, Wang L, Liu J, Zhang Z, Tang Y, et al. The Cotton miR477-CBP60A Module Participates in Plant Defense Against Verticillium dahlia. Mol Plant Microbe Interact 2020;33:624–36. 10.1094/MPMI-10-19-0302-R.

[6] Salvador-Guirao R, Baldrich P, Weigel D, Rubio-Somoza I, San Segundo B. The MicroRNA miR773 Is Involved in the Arabidopsis Immune Response to Fungal Pathogens. Mol Plant Microbe Interact 2018;31:249–59. 10.1094/MPMI-05-17-0108-R.

[7] Thiebaut F, Rojas CA, Grativol C, Motta MR, Vieira T, Regulski M, et al. Genome-wide identification of microRNA and siRNA responsive to endophytic beneficial diazotrophic bacteria in maize. BMC Genomics 2014;15:766. 10.1186/1471-2164-15-766.

[8] Ben Amor B, Wirth S, Merchan F, Laporte P, d’Aubenton-Carafa Y, Hirsch J, et al. Novel long non-protein coding RNAs involved in Arabidopsis differentiation and stress responses. Genome Res 2009;19:57–69. 10.1101/gr.080275.108.

[9] Lin X, Lin W, Ku Y-S, Wong F-L, Li M-W, Lam H-M, et al. Analysis of Soybean Long Non-Coding RNAs Reveals a Subset of Small Peptide-Coding Transcripts. Plant Physiol 2020;182:1359–74. 10.1104/pp.19.01324.

[10] Gloss BS, Dinger ME. The specificity of long noncoding RNA expression. Biochim Biophys Acta 2016;1859:16–22. 10.1016/j.bbagrm.2015.08.005.

[11] Zhang Y-C, Liao J-Y, Li Z-Y, Yu Y, Zhang J-P, Li Q-F, et al. Genome-wide screening and functional analysis identify a large number of long noncoding RNAs involved in the sexual reproduction of rice. Genome Biol 2014;15:512. 10.1186/s13059-014-0512-1.

[12] Li L, Eichten SR, Shimizu R, Petsch K, Yeh C-T, Wu W, et al. Genome-wide discovery and characterization of maize long non-coding RNAs. Genome Biol 2014;15:R40. 10.1186/gb-2014-15-2-r40.

[13] Ward M, McEwan C, Mills JD, Janitz M. Conservation and tissue-specific transcription patterns of long noncoding RNAs. Journal of Human Transcriptome 2015;1:2–9. 10.3109/23324015.2015.1077591.

[14] Wang KC, Yang YW, Liu B, Sanyal A, Corces-Zimmerman R, Chen Y, et al. A long noncoding RNA maintains active chromatin to coordinate homeotic gene expression. Nature 2011;472:120–4. 10.1038/nature09819.

[15] Heo JB, Sung S. Vernalization-mediated epigenetic silencing by a long intronic noncoding RNA. Science 2011;331:76–9. 10.1126/science.1197349.

[16] Guil S, Esteller M. Cis-acting noncoding RNAs: friends and foes. Nat Struct Mol Biol 2012;19:1068–75. 10.1038/nsmb.2428.

[17] Urquiaga MC de O, Thiebaut F, Hemerly AS, Ferreira PCG. From trash to luxury: the potential role of plant lncrna in DNA methylation during abiotic stress. Front Plant Sci 2020;11:603246. 10.3389/fpls.2020.603246.

[18] Wang Y, Luo X, Sun F, Hu J, Zha X, Su W, et al. Overexpressing lncRNA LAIR increases grain yield and regulates neighbouring gene cluster expression in rice. Nat Commun 2018;9:3516. 10.1038/s41467-018-05829-7.

[19] Fang J, Zhang F, Wang H, Wang W, Zhao F, Li Z, et al. Ef-cd locus shortens rice maturity duration without yield penalty. Proc Natl Acad Sci USA 2019;116:18717–22. 10.1073/pnas.1815030116.

[20] Wang A, Hu J, Gao C, Chen G, Wang B, Lin C, et al. Genome-wide analysis of long non-coding RNAs unveils the regulatory roles in the heat tolerance of Chinese cabbage (Brassica rapa ssp.chinensis). Sci Rep 2019;9:5002. 10.1038/s41598-019-41428-2.

[21] Wang T-Z, Liu M, Zhao M-G, Chen R, Zhang W-H. Identification and characterization of long non-coding RNAs involved in osmotic and salt stress in Medicago truncatula using genome-wide high-throughput sequencing. BMC Plant Biol 2015;15:131. 10.1186/s12870-015-0530-5.

[22] Zhang W, Han Z, Guo Q, Liu Y, Zheng Y, Wu F, et al. Identification of maize long non-coding RNAs responsive to drought stress. PLoS ONE 2014;9:e98958. 10.1371/journal.pone.0098958.

[23] Chen J, Zhong Y, Qi X. LncRNA TCONS_00021861 is functionally associated with drought tolerance in rice (Oryza sativa L.) via competing endogenous RNA regulation. BMC Plant Biol 2021;21:410. 10.1186/s12870-021-03195-z.

[24] Golicz AA, Singh MB, Bhalla PL. The long intergenic noncoding RNA (lincrna) landscape of the soybean genome. Plant Physiol 2018;176:2133–47. 10.1104/pp.17.01657.

[25] Golicz AA, Bhalla PL, Singh MB. MCRiceRepGP: a framework for the identification of genes associated with sexual reproduction in rice. Plant J 2018;96:188–202. 10.1111/tpj.14019.

[26] Zhu Q-H, Stephen S, Taylor J, Helliwell CA, Wang M-B. Long noncoding RNAs responsive to Fusarium oxysporum infection in Arabidopsis thaliana. New Phytol 2014;201:574–84. 10.1111/nph.12537.

[27] Zhang H, Hu W, Hao J, Lv S, Wang C, Tong W, et al. Genome-wide identification and functional prediction of novel and fungi-responsive lincRNAs in Triticum aestivum. BMC Genomics 2016;17:238. 10.1186/s12864-016-2570-0.

[28] Xin M, Wang Y, Yao Y, Song N, Hu Z, Qin D, et al. Identification and characterization of wheat long non-protein coding RNAs responsive to powdery mildew infection and heat stress by using microarray analysis and SBS sequencing. BMC Plant Biol 2011;11:61. 10.1186/1471-2229-11-61.

[29] Kang Y-J, Yang D-C, Kong L, Hou M, Meng Y-Q, Wei L, et al. CPC2: a fast and accurate coding potential calculator based on sequence intrinsic features. Nucleic Acids Res 2017;45:W12–6. 10.1093/nar/gkx428.

[30] Cagirici HB, Galvez S, Sen TZ, Budak H. LncMachine: a machine learning algorithm for long noncoding RNA annotation in plants. Funct Integr Genomics 2021;21:195–204. 10.1007/s10142-021-00769-w.

[31] Wang L, Park HJ, Dasari S, Wang S, Kocher J-P, Li W. CPAT: Coding-Potential Assessment Tool using an alignment-free logistic regression model. Nucleic Acids Res 2013;41:e74. 10.1093/nar/gkt006.

[32] Singh U, Khemka N, Rajkumar MS, Garg R, Jain M. PLncPRO for prediction of long non-coding RNAs (lncRNAs) in plants and its application for discovery of abiotic stress-responsive lncRNAs in rice and chickpea. Nucleic Acids Res 2017;45:e183. 10.1093/nar/gkx866.

[33] Li A, Zhang J, Zhou Z. PLEK: a tool for predicting long non-coding RNAs and messenger RNAs based on an improved k-mer scheme. BMC Bioinformatics 2014;15:311. 10.1186/1471-2105-15-311.

[34] Li J, Zhang X, Liu C. The computational approaches of lncRNA identification based on coding potential: Status quo and challenges. Comput Struct Biotechnol J 2020;18:3666–77. 10.1016/j.csbj.2020.11.030.

[35] Pian C, Zhang G, Chen Z, Chen Y, Zhang J, Yang T, et al. LncRNApred: Classification of Long Non-Coding RNAs and Protein-Coding Transcripts by the Ensemble Algorithm with a New Hybrid Feature. PLoS ONE 2016;11:e0154567. 10.1371/journal.pone.0154567.

[36] Fan X-N, Zhang S-W. lncRNA-MFDL: identification of human long non-coding RNAs by fusing multiple features and using deep learning. Mol Biosyst 2015;11:892–7. 10.1039/c4mb00650j.

[37] Eraslan G, Avsec Ž, Gagneur J, Theis FJ. Deep learning: new computational modelling techniques for genomics. Nat Rev Genet 2019;20:389–403. 10.1038/s41576-019-0122-6.

[38] Shrestha A, Mahmood A. Review of deep learning algorithms and architectures. IEEE Access 2019;7:53040–65. 10.1109/ACCESS.2019.2912200.

[39] Szcześniak MW, Bryzghalov O, Ciomborowska-Basheer J, Makałowska I. Cantatadb 2.0: expanding the collection of plant long noncoding rnas. Methods Mol Biol 2019;1933:415–29. 10.1007/978-1-4939-9045-0_26.

[40] Jin J, Lu P, Xu Y, Li Z, Yu S, Liu J, et al. PLncDB V2.0: a comprehensive encyclopedia of plant long noncoding RNAs. Nucleic Acids Res 2020;49:D1489–95. 10.1093/nar/gkaa910.

[41] Paytuví Gallart A, Hermoso Pulido A, Anzar Martínez de Lagrán I, Sanseverino W, Aiese Cigliano R. GREENC: a Wiki-based database of plant lncRNAs. Nucleic Acids Res 2016;44:D1161–6. 10.1093/nar/gkv1215.

[42] Di Marsico M, Paytuvi Gallart A, Sanseverino W, Aiese Cigliano R. GreeNC 2.0: a comprehensive database of plant long non-coding RNAs. Nucleic Acids Res 2021. 10.1093/nar/gkab1014.

[43] Singh A, Vivek AT, Kumar S. AlnC: An extensive database of long non-coding RNAs in angiosperms. PLoS ONE 2021;16:e0247215. 10.1371/journal.pone.0247215.

[44] Devlin J, Chang M-W, Lee K, Toutanova K. Bert: Pre-training of deep bidirectional transformers for language understanding. ArXiv Preprint ArXiv:181004805 2018.

[45] Howard J, Ruder S. Universal language model fine-tuning for text classification. ArXiv Preprint ArXiv:180106146 2018.

[46] Vaswani A, Shazeer N, Parmar N, Uszkoreit J, Jones L, Gomez AN, et al. Attention Is All You Need. ArXiv 2017:5998–6008.

[47] Ji Y, Zhou Z, Liu H, Davuluri RV. DNABERT: pre-trained Bidirectional Encoder Representations from Transformers model for DNA-language in genome. Bioinformatics 2021;37:2112–20. 10.1093/bioinformatics/btab083.

[48] Mo S, Fu X, Hong C, Chen Y, Zheng Y, Tang X, et al. Multi-modal Self-supervised Pre-training for Regulatory Genome Across Cell Types. ArXiv 2021.

[49] Wahab A, Tayara H, Xuan Z, Chong KT. DNA sequences performs as natural language processing by exploiting deep learning algorithm for the identification of N4-methylcytosine. Sci Rep 2021;11:212. 10.1038/s41598-020-80430-x.

[50] Meng X, Liang Z, Dai X, Zhang Y, Mahboub S, Ngu DW, et al. Predicting transcriptional responses to cold stress across plant species. Proc Natl Acad Sci USA 2021;118. 10.1073/pnas.2026330118.

[51] Pierse C. Transformers interpret. Model Explainability That Works Seamlessly with Transformers 2021. https://github.com/cdpierse/transformers-interpret (accessed January 21, 2022).

[52] Fico A, Fiorenzano A, Pascale E, Patriarca EJ, Minchiotti G. Long non-coding RNA in stem cell pluripotency and lineage commitment: functions and evolutionary conservation. Cell Mol Life Sci 2019;76:1459–71. 10.1007/s00018-018-3000-z.

[53] Jiao Y, Peluso P, Shi J, Liang T, Stitzer MC, Wang B, et al. Improved maize reference genome with single-molecule technologies. Nature 2017;546:524–7. 10.1038/nature22971.

[54] Rošić S, Erhardt S. No longer a nuisance: long non-coding RNAs join CENP-A in epigenetic centromere regulation. Cell Mol Life Sci 2016;73:1387–98. 10.1007/s00018-015-2124-7.

[55] Szcześniak MW, Bryzghalov O, Ciomborowska-Basheer J, Maka\lowska I. CANTATAdb 2.0: expanding the collection of plant long noncoding RNAs. Plant Long Non-Coding RNAs, Humana Press, New York, NY; 2019, p. 415–429.

[56] Barbedo JGA. Impact of dataset size and variety on the effectiveness of deep learning and transfer learning for plant disease classification. Computers and Electronics in Agriculture 2018;153:46–53. 10.1016/j.compag.2018.08.013.

[57] Wang X, Wang H, Wang J, Sun R, Wu J, Liu S, et al. The genome of the mesopolyploid crop species Brassica rapa. Nat Genet 2011;43:1035–9. 10.1038/ng.919.

[58] Kawahara Y, de la Bastide M, Hamilton JP, Kanamori H, McCombie WR, Ouyang S, et al. Improvement of the *Oryza sativa* Nipponbare reference genome using next generation sequence and optical map data. Rice (N Y) 2013;6:4. 10.1186/1939-8433-6-4.

[59] Johnsson P, Lipovich L, Grandér D, Morris KV. Evolutionary conservation of long non-coding RNAs; sequence, structure, function. Biochim Biophys Acta 2014;1840:1063–71. 10.1016/j.bbagen.2013.10.035.

[60] Deng P, Liu S, Nie X, Weining S, Wu L. Conservation analysis of long non-coding RNAs in plants. Sci China Life Sci 2018;61:190–8. 10.1007/s11427-017-9174-9.

[61] Tavares RCA, Pyle AM, Somarowthu S. Phylogenetic Analysis with Improved Parameters Reveals Conservation in lncRNA Structures. J Mol Biol 2019;431:1592–603. 10.1016/j.jmb.2019.03.012.

[62] Hezroni H, Koppstein D, Schwartz MG, Avrutin A, Bartel DP, Ulitsky I. Principles of long noncoding RNA evolution derived from direct comparison of transcriptomes in 17 species. Cell Rep 2015;11:1110–22. 10.1016/j.celrep.2015.04.023.

[63] RNAcentral Consortium. RNAcentral 2021: secondary structure integration, improved sequence search and new member databases. Nucleic Acids Res 2021;49:D212–20. 10.1093/nar/gkaa921.

[64] The RNAcentral Consortium, Petrov AI, Kay SJE, Kalvari I, Howe KL, Gray KA, et al. RNAcentral: a comprehensive database of non-coding RNA sequences. Nucleic Acids Res 2017;45:D128–34. 10.1093/nar/gkw1008.

[65] Pruitt KD, Tatusova T, Maglott DR. NCBI Reference Sequence (RefSeq): a curated non-redundant sequence database of genomes, transcripts and proteins. Nucleic Acids Res 2005;33:D501–4. 10.1093/nar/gki025.

[66] Zhou B, Ji B, Liu K, Hu G, Wang F, Chen Q, et al. EVLncRNAs 2.0: an updated database of manually curated functional long non-coding RNAs validated by low-throughput experiments. Nucleic Acids Res 2021;49:D86–91. 10.1093/nar/gkaa1076.

[67] Quinlan AR, Hall IM. BEDTools: a flexible suite of utilities for comparing genomic features. Bioinformatics 2010;26:841–2. 10.1093/bioinformatics/btq033.

[68] Camacho C, Coulouris G, Avagyan V, Ma N, Papadopoulos J, Bealer K, et al. BLAST+: architecture and applications. BMC Bioinformatics 2009;10:421. 10.1186/1471-2105-10-421.

[69] Li H, Handsaker B, Wysoker A, Fennell T, Ruan J, Homer N, et al. The Sequence Alignment/Map format and SAMtools. Bioinformatics 2009;25:2078–9. 10.1093/bioinformatics/btp352.

[70] Kurtzer GM, Sochat V, Bauer MW. Singularity: Scientific containers for mobility of compute. PLoS ONE 2017;12:e0177459. 10.1371/journal.pone.0177459.

[71] Fabian P, Gael V, Alexandre G, Vincent M, Bertrand T. Scikit-learn: Machine Learning in Python. J Mach Learn Res 2011;12:2825--2830.

[72] Wickham H, Averick M, Bryan J, Chang W, McGowan L, François R, et al. Welcome to the tidyverse. JOSS 2019;4:1686. 10.21105/joss.01686.

[73] Wickham H. ggplot2. WIREs Comp Stat 2011;3:180–5. 10.1002/wics.147.

