## Supplementary figures and images for "DNABERT-based explainable lncRNA identification in plant genome assemblies"

### Supplementary Figures

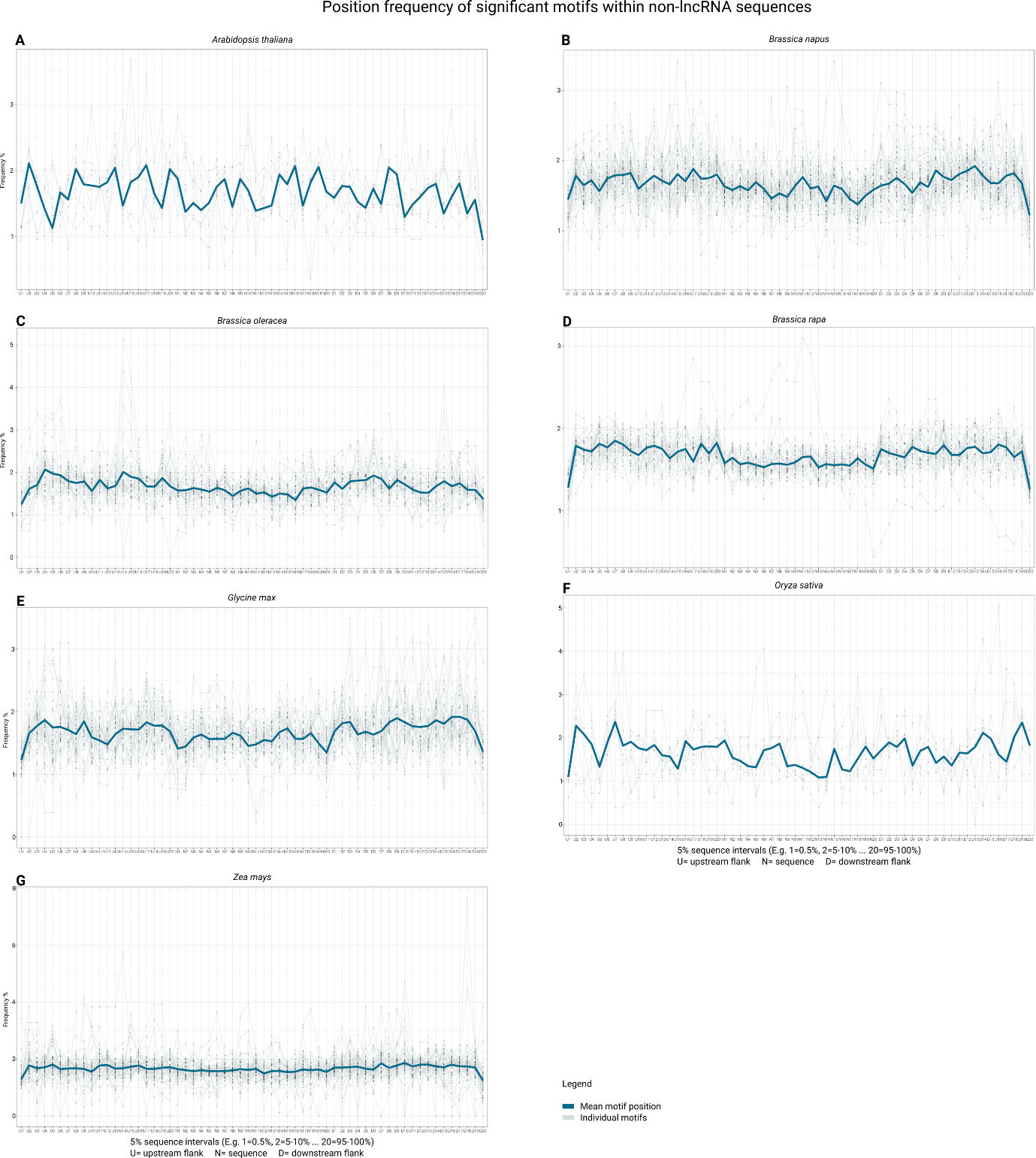

Supplementary Figure 1: Position frequency of significant motifs within non-lncRNA sequences
